# Interaction between host genes and M. tuberculosis lineage can affect tuberculosis severity: evidence for coevolution

**DOI:** 10.1101/769448

**Authors:** Michael L. McHenry, Jacquelaine Bartlett, Robert P. Igo, Edward Wampande, Penelope Benchek, Harriet Mayanja-Kizza, Kyle Fluegge, Noemi B. Hall, Sebastien Gagneux, Sarah A. Tishkoff, Christian Wejse, Giorgio Sirugo, W. Henry Boom, Moses Joloba, Scott M. Williams, Catherine M. Stein

## Abstract

Genetic studies of both the human host and *Mycobacterium tuberculosis* (MTB) demonstrate independent association with tuberculosis (TB) risk. However, neither explains a large portion of disease risk or severity. Based on studies in other infectious diseases and animal models of TB, we hypothesized that the genomes of the two interact to modulate risk of developing active TB or increasing the severity of disease, when present. We examined this hypothesis in our TB household contact study in Kampala, Uganda, in which there were 3 MTB lineages of which L4-Ugandan (L4.6) is the most recent. TB severity, measured using the Bandim TBscore, was modeled as a function of host SNP genotype, MTB lineage, and their interaction, within two independent cohorts of TB cases, N=113 and 122. No association was found between lineage and severity, but association between multiple polymorphisms in IL12B and TBscore was replicated in two independent cohorts (most significant rs3212227, combined p=0.0006), supporting previous associations of IL12B with TB susceptibility. We also observed significant interaction between a single nucleotide polymorphism (SNP) in SLC11A1 and the L4-Ugandan lineage in both cohorts (rs17235409, meta p=0.0002). Interestingly, the presence of the L4-Uganda lineage in the presence of the ancestral human allele associated with more severe disease. These findings demonstrate that IL12B is associated with severity of TB in addition to susceptibility, and that the association between TB severity and human genetics can be due to an interaction between genes in the two species, providing evidence of host-pathogen coevolution in TB.

**AUTHOR SUMMARY:** Susceptibility to tuberculosis (TB) is affected by genetic variation in both the human host and the causative bacterium, *Mycobacterium tuberculosis*. However, prior studies of the genetics of each species have not explained a large part of TB risk. The possibility exists that risk can be better estimated from patterns of variation in two species as a unit, such that some combinations provide increased risk, or in the presence of TB, increased disease severity. We hypothesized that alleles in the two species that have co-existed for long periods are more likely to reduce disease severity so as to promote prolonged co-occurrence. We tested this by studying TB severity in two patient cohorts from Uganda for which paired MTB-human DNA were available. We examined severity, as measured by the Bandim TBscore, and assessed whether there was an interaction between MTB lineage and SNPs in the host with this metric. Our results indicate that the most recent TB lineage (L4.6/Uganda) when found together with an ancestral allele in SLC11A1 resulted in more severe disease. This finding is consistent with the conclusion that MTB and human have coevolved to modulate TB severity.

## Introduction

Pulmonary tuberculosis (TB), a respiratory disease caused by *Mycobacterium tuberculosis* (MTB) infection, creates a significant public health burden worldwide, with 10 million incident cases and an estimated 1.64 million deaths in 2017[1]. Susceptibility to pulmonary TB can be influenced by human genetic variation with both candidate gene and genome-wide studies having identified variants that affect risk of disease[2-8]. However, to our knowledge there has been only one study characterizing TB severity as a quantitative trait and examining genetic associations with this trait[9].

There is evidence that MTB genetic variation as delineated by phylogenetic lineage can independently affect disease sequelae[10-13]. Recent work has classified MTB into at least seven genetic lineages that are defined both by geography and a historical timeline of human exposure [14]. There are also specialized sub-lineages that are more recently divergent and restricted to smaller geographic areas, including one found solely in Uganda and its neighboring countries known as the L4.6/Uganda sub-lineage[15-18]. Additionally, prior evidence supports these independent roles of both host and MTB genetic variation on disease risk or outcomes, but it is unclear whether interactions between the genomes of the two species influence disease. Studies examining the interaction between host genotype and MTB lineage are sparse, especially those examining severity as an outcome.

Long-term coexistence between the human genome and MTB lineage may decrease risk of developing active TB or minimize the severity of disease, if present [19]. Specifically, coevolution implies distinct historical and geographic variation in the prevalence of MTB lineages that would allow the two species to adapt to each other, thereby enabling coordinated evolution between host and MTB genotypes. If such interactions exist it should be possible to measure disease risk or severity in terms of human–MTB coevolution as a result of historical coexistence[20]. If this is the case, it would represent an example of prudent exploitation or covert infection in which infection does not necessarily lead to active disease and may cause less virulent disease, when present[21, 22]. In fact, most people exposed to MTB do not progress to active disease[23]. Under the coevolution model, a newly divergent MTB lineage (that has not historically co-existed with the population in question) is expected to cause more severe disease[24]. The potential for human-MTB coevolution has been explored in human and model systems, but studies have not yet identified a definitive effect at the population level[3, 4, 10, 13, 25-27]. Although coevolution may affect the likelihood of developing active disease, to study coevolution in practice, it is necessary to study cases only, as we cannot deduce the MTB lineage(s) to which unaffected individuals have been exposed with certainty. Even within a household, the strain to which an individual is predominantly exposed may not match that of the index case, since community exposure can be the major source of exposure[28].

Because unaffected individuals cannot be examined, all previous studies of host-MTB genome interaction have been case-only studies that at best examine association between lineage and host genotypes, but not explicit interactions [25, 26, 29, 30]. Host-pathogen coevolution has been demonstrated in other organisms, by removing an apparent independent effect of host genetic factors on disease severity, thereby adjusting for host-pathogen interactions[19, 31]. In TB, this has been problematic, until the recent development of the Bandim TBscore that provides a novel means to objectively assess TB severity[32].The Bandim score can predict mortality and associates with procalcitonin and C-reactive protein, biomarkers of TB severity[33-36]. In the present study, this measure was used to assess severity of TB and the role of coevolution between MTB and humans. A distinct advantage of the TBScore is that it is based on simple measures of clinical parameters that can be determined in resource-limited environments where TB is most prevalent.

In this study of genes implicated in TB pathogenesis, we assessed the role of human genetic variation and MTB lineage in TB severity, and whether the two interact. We used a case-only approach to unambiguously identify MTB lineages in exposed individuals. This approach will help us elucidate the degree to which MTB is a prudent parasite whose evolution has mirrored that of its human host. We are unaware of any previous studies that have examined the role of coevolution on TB severity or the role of human genetic variation on TB severity as a quantitative trait, making this study the first of its kind.

## Results

### Population Characteristics

Of the 113 subjects in cohort 1, 51 (45.2%) were female, and 46 (40.7%) were HIV positive (Table 1). Subjects ranged from age 16 to 67 and the mean age was 29.74 with a standard deviation of 9.1 years. The mean TBscore was 5.86 with a standard deviation of 2.20. Of the 121 subjects in cohort 2, 51 (42.2%) were female, and 31 (25.6%) were HIV positive. Subjects ranged in age from 15 to 70 with a mean of 29.17 and a standard deviation of 8.9. The mean TBscore was 5.53 with standard deviation of 2.21. The two cohorts were similar in every aspect except for HIV positive status (Table 1). Thus, as stated in our statistical methods section, we controlled for HIV status in all of our regression models so that HIV status would not alter our conclusions.

**Table 1:** Cohort Demographics and Clinical Measures.

| Variable | Total | Cohort 1 | Cohort 2 | p* |
| --- | --- | --- | --- | --- |
| Sample Size | 234 | 113 | 121 | - |
| Female | 102 (43.6%) | 51(45.2%) | 51(42.2%) | 0.65 |
| HIV+ | 77(32.9%) | 46 (40.7%) | 31(25.6%) | 0.014 |
| Age, years | 29.44 ± 8.9 | 29.74 ± 9.1 | 29.17 ± 8.9 | 0.62 |
| Age range, years | 15-70 | 16-67 | 15-70 | - |
| TBscore | 5.69 ± 2.21 | 5.86 ± 2.20 | 5.53 ± 2.21 | 0.25 |

Values are shown as N (%) or as mean ± SD. \*Comparisons between Cohort 1 and Cohort 2 were made using Pearson's $\chi^2$ test and for continuous variables using Student's t-test.

### Lineage and Severity

There were 3 lineages in this study: L3: Central Asia, L4/Non-Ugandan, and L4.6/Uganda. Of these, the specialist L4.6/Uganda sub-lineage is thought to be the most recently divergent and is only found in Uganda and its neighboring countries. The L3 and L4 generalist lineages have a longer history and are more widespread[14-18]. Severity of TB was not associated with lineage independently of human genotype (Figure 1, p=0.71 by ANOVA), consistent with previous findings with clinical characteristics considered singly[18]. Comparisons of only the L4.6/Uganda to the other two lineages also indicated no differences in disease severity (p=0.44). In addition, lineage distribution did not differ significantly between cohorts (p = 0.58; Supplemental Table 1).

**Figure 1:**
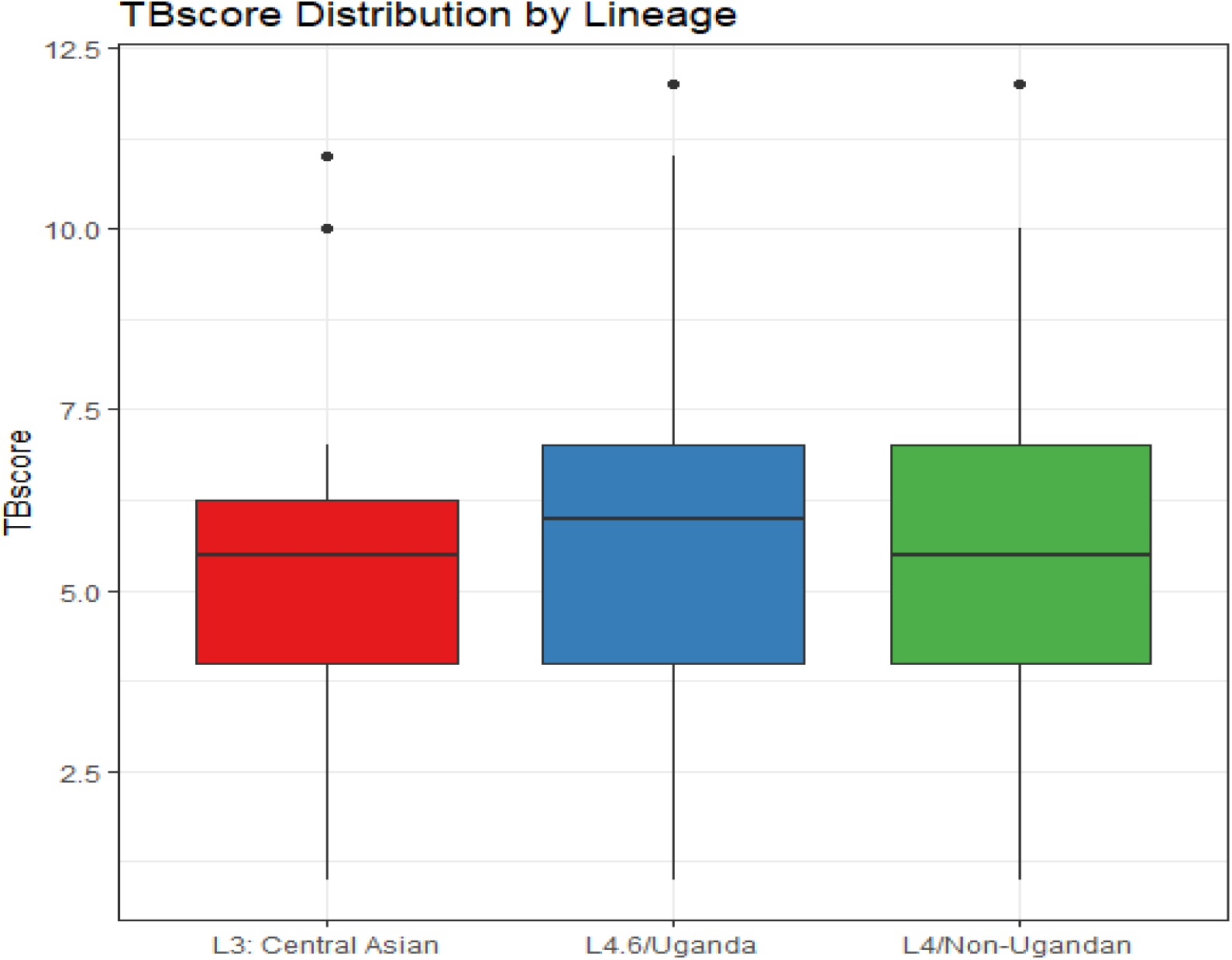
TBscore across Lineages. Lines within each lineage show the median values. Mean TBscore (+/-S.D.) is 5.6 (+/-2.2) for L4/Non-Ugandan; 5.4 (+/-2.4) for L3:Central Asia and 5.8 (+/-2.2) for L4.6/Uganda. These do not differ p = 0.71 (ANOVA).

### Independent effects of MTB lineage and Human SNPs on TB score

Of the 403 SNPs tested, we observed significant association between multiple SNPs in *IL12B* and TBscore that was consistent across cohorts. Three SNPs were nominally significant (p < 0.05) in Cohort 1 and all replicated (p < 0.05) in Cohort 2. A fourth SNP (rs3213094) was borderline significant in Cohort 1 (p = 0.06) but was significant in Cohort 2 and the combined data (p = 0.0009). The three SNPs that were significant in both sets showed perfect linkage disequilibrium (LD) (r^2^ ≥0.99 in all pairwise comparisons for the combined data (Table 2 and Supplemental Figures 1-3); rs3213094 showed similarly high levels of LD with the other SNPs (Supplemental Figures 1-2). The SNP with the most significant combined p-value was rs3212220 (p=0.0005). All of these SNPs had similar effect sizes (β values) in the linear regression model, ranging from 0.98 to 1.00. The presence of the homozygous ancestral genotype was associated with a one-point average increase in the TBscore, equivalent to one more clinically relevant symptom, meaning the derived allele is associated with lower severity. As a score of 3 or greater is a predictor of 18 month mortality; therefore, a one point increase is clinically meaningful in some contexts[32]. While association with SNPs in other genes, *NOD1* and *STAT1*, was observed, strict replication was not obtained, as the same SNPs were not nominally significant (p < 0.05) across both samples (Supplemental Table 2). Different SNPs in these genes were significant in at least one of the cohorts.

**Table 2:** Association between *IL12B*Markers and TBscore, adjusted for HIV Status.

| Name | Alleles | Notation | Cohort 1 |  | Cohort 2 |  | Combined |  |
| --- | --- | --- | --- | --- | --- | --- | --- | --- |
| | | | $\beta$ (95% CI) | p | $\beta$ (95% CI) | p | $\beta$ (95% CI) | p |
| rs3212227 | **T/G** | 3' UTR | 0.94 (0.08, 1.79) | 0.03 | 1.05 (0.25, 1.85) | 0.01 | 1.00 (0.44, 1.58) | 0.0006 |
| rs3212219 | **C/A** | Intron | 0.89 (0.04, 1.74) | 0.04 | 1.05 (0.25, 1.86) | 0.01 | 0.98 (0.43, 1.56) | 0.0010 |
| rs3212220 | **C/A** | Intron | 0.91 (0.01, 1.81) | 0.05 | 1.05 (0.25, 1.87) | 0.01 | 1.01 (0.40, 1.56) | 0.0005 |
| rs3213094 | **C/T** | Intron | 0.84 (−0.02, 1.71) | 0.06 | 1.05 (0.25, 1.85) | 0.01 | 0.98 (0.41, 1.55) | 0.0009 |

All analyses coded using homozygous ancestral as 1 and one or more copies of the derived allele coded as 0. The ancestral allele is bold in the table.

### Examination of host-pathogen coevolution

The interaction between SNP and lineage was assessed with a linear regression model that included a multiplicative term for SNP and lineage interactions (Supplemental Table 3). Interactions were considered significant only if p < 0.05 for the interaction term in both cohorts. Our analyses were focused on contrast between L4.6/Uganda and the other two lineages combined. This strategy also permitted us to assess the role of the derived L4.6/Ugandan lineage, with which Ugandans have spent less time, in TB severity. A significant interaction between one SNP in *SLC11A1*, rs17235409, and L4.6/Uganda was observed in both cohorts. The effect for this interaction was in the same direction and of similar value in each cohort and showed a more significant association in the combined analysis (combined p=0.00022) (Table 3), significant even after Bonferroni correction (see Supplementary methods). Of note, rs17235409 is a non-synonymous exonic variant (D543N). Furthermore, this SNP was not associated with TBscore in the absence of the interaction term with lineage. The effect size (β value) for the interaction in the combined analysis was 2.63. To interpret this, consider that there are four possible combinations of genotype and lineage in this analysis, holding HIV status constant. The combinations with the highest TBscore were: (1) the carriers of derived alleles (AA, GA) with the lineages that are more widespread and have had longer historical contact with humans (L3 and L4) and (2) the ancestral homozygous genotype (GG) with the more newly evolved Ugandan sub-lineage (L4.6). This is important as we observe the lowest average TBscore for the combination of ancestral allele and older lineages, i.e. the genotype and lineage that have historically co-existed. We observe the highest average TBscore for the combinations of genotype and lineage that have not historically co-existed. These findings support our hypothesis that coevolution between a lineage and genotype would associate with lower severity. For this SNP, the allele that associates with more severe disease in the interaction model is the ancestral human allele, G (Figure 2; Table 3). In both cohorts, the simultaneous presence of both the ancestral human allele and the derived L4.6/Uganda sub-lineage was associated with increased TBscore, indicating that evolutionary histories of both species, taken together, affected disease severity. That these associate only in the presence of an interaction term is indicative of coevolution between human variants and MTB lineage.

**Table 3:** Full Regression model results for*SLC11A1* Marker rs17235409.

|  | Cohort 1 |  | Cohort 2 |  | Combined |  |
| --- | --- | --- | --- | --- | --- | --- |
| | $\beta$ (95% CI) | p | $\beta$ (95% CI) | p | $\beta$ (95% CI) | p |
| rs17235409( $\beta_3$)† | -2.53 (-4.29, -0.77) | 0.0057 | -1.85 (-3.29, -0.40) | 0.014 | -2.11 (-1.00, -3.23) | 0.00026 |
| L4.6/Ugandan($\beta_2$) | -1.43 (-3.36, 0.50) | 0.15 | -2.18 (-3.73, -0.63) | 0.0067 | -1.83 (-0.62, -3.03) | 0.00331 |
| HIV+ Status($\beta_1$) | 0.30 (-0.51, 1.11) | 0.47 | -0.47 (-1.39, 0.44) | 0.31 | 0.033 (-0.56, 0.63) | 0.911 |
| rs17235409*L4.6/Ugandan‡ (combination of GG and L4.6/Ugandan) ($\beta_4$) | 2.36 (0.21, 4.50) | 0.033 | 2.71(0.91, 4.51) | 0.0039 | 2.63 (1.25, 4.00) | 0.00022 |

rs17235409 is an exonic SNP at position 219259732 on chromosome 2. The ancestral allele is G and the derived is A.
Regression Model:
$X_1$ = rs17235409, $X_2$ =HIV+ Status, $X_3$ = Ugandan Lineage
†Model of inheritance (GG vs GA/AA as referent)
‡Combination of GA/AA and L4-Ugandan) ( $\beta_4$ )

**Figure 2:**
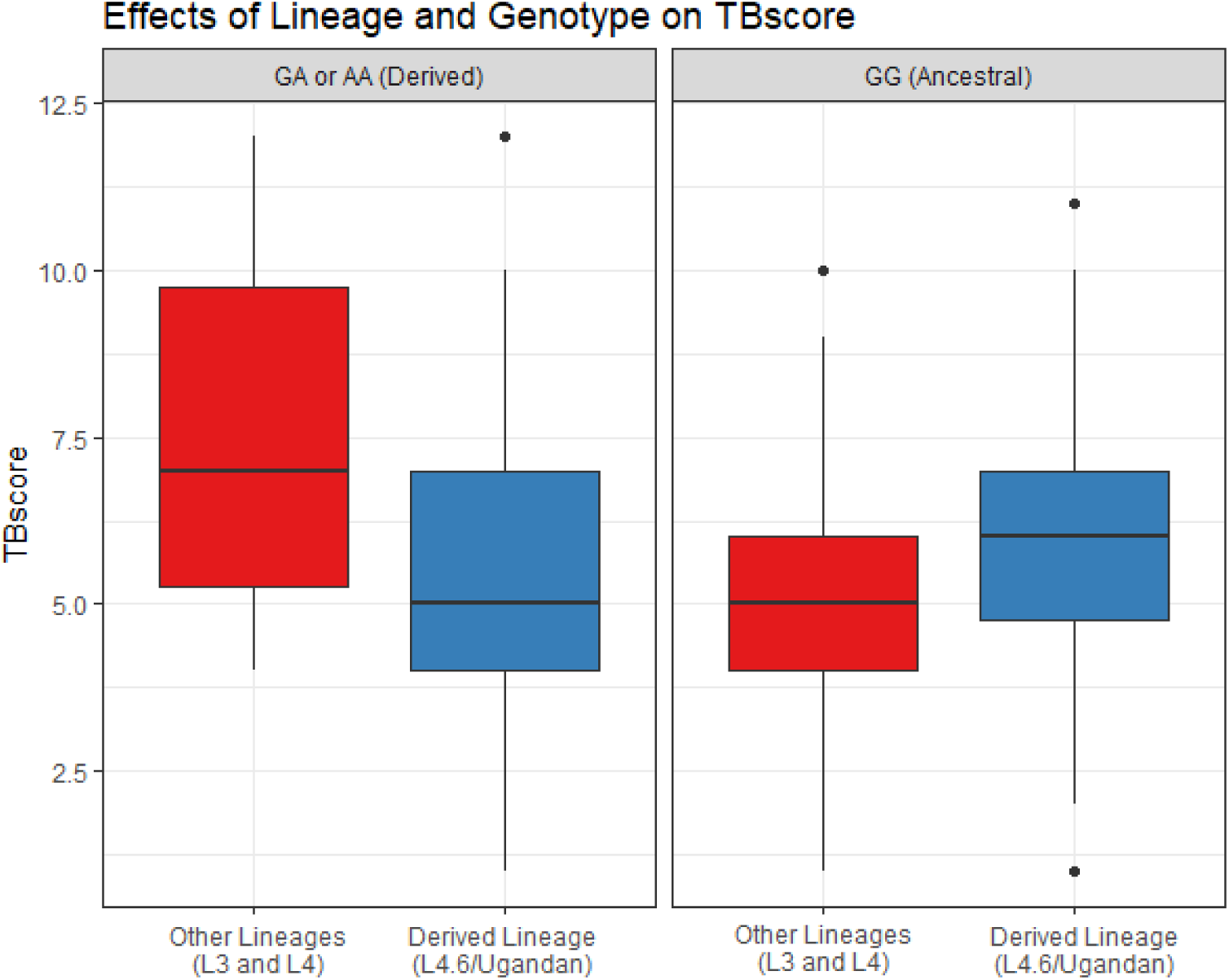
Effects of rs17235409 (SLC11A1) Genotype and Lineage on TBscore. The association of TB severity with lineage differs by genotype. TBscore is smaller in the Uganda lineage for individuals carrying a derived allele (GA or AA)(left panel) at *SLC11A1* but larger for the ancestral genotype (GG) carriers (right panel). These results represent a significant interaction between rs17235409 and MTB lineage (p= 0.00022).

## Discussion

Our hypothesis is that coevolution between humans and *M. tuberculosis* in the same area may lead to less severe disease when host individuals possess the ancestral allele but are infected with a historically co-existing lineage of MTB [37]. Our observation of more severe TB when the human homozygous ancestral genotype (GG) was combined with the more recently divergent L4.6/Uganda MTB sub-lineage and less severe TB when this younger lineage is combined with a derived genotype is in line with the conclusion that disrupted coevolution increases disease severity. The L4.6/Uganda lineage, almost exclusively found in Uganda and neighboring countries, is the most recently diverged clade in our sample, and is therefore not the lineage Ugandans have historically been infected with. Therefore, we posit that individuals with ancestral Ugandan human genotypes may not be able to mount as effective an immune response as they have not historically been exposed to L4.6/Uganda, the derived lineage. Additionally, if an MTB lineage has evolved in the context of specific human standing variation, then we would hypothesize a derived lineage to cause more severe disease, as we observed in Uganda, consistent with the hypothesis of prudent exploitation.

In our two independent sets of subjects, we identified both marginal effects for TB severity in loci previously associated with TB disease (*IL12B*) and evidence for interactions between human SNPs in *SLC11A1* and MTB lineage. Of note, the *SLC11A1* SNP that interacted with MTB lineage did not associate in the absence of interaction with TBscore. These interactions in the absence of marginal effects have previously been considered a hallmark of coevolution[19, 31].

The genes we have analyzed in both the marginal analysis and interaction analysis were chosen because they are all biologically plausible as they produce end-products known to be important in the immune response to MTB and have been previously identified as modifiers of responses to MTB infection[3, 38]. *IL12B* is an important regulator of immune responses to MTB, including IFN-γ secretion. A previous epidemiologic study has suggested *IL12B* as a determinant of TB susceptibility in humans, and previous studies have shown associations with one of the same SNPs that we replicated in both sets (rs3212227)[8, 39]. Additionally, prior work has shown that IL12 pathway deficiency associates with mycobacterial diseases[40]. Lastly, murine models have shown that an IL12 gene knockout renders mice highly susceptible to MTB infection[41]. While *IL12B* has previously been shown to be important in TB, these studies did not examine severity. We believe that previous evidence in the context of susceptibility is consistent with our findings that *IL12B* is an important determinant of TB disease severity but it is important to consider the possibility that distinct biological processes are underlying each phenotype and thus this result is a novel finding that adds to previous literature about the putative role of *IL12B* in TB pathogenesis. Further, very few studies have examined clinical severity as a phenotype and it is important to expand this area of study.

*SLC11A1* (previously called *NRAMP1*), an important regulator of macrophage responses to MTB, has long been implicated in resistance to intracellular infections, and may have a role in mycobacterial susceptibility at a population level[5, 42, 43]. *SLC11A1* has been well characterized and is thought to code for a membrane-bound divalent cation transporter found exclusively in macrophages and polymorphonuclear cells that has pleiotropic effects on macrophage activation. It functions on the phagolyosomal surface in macrophages, regulating changes in iron transport and the transport of other cations in response to infection. Further, MTB expresses a membrane-bound cation transporter in the *SLC11* family that functions similarly to, and potentially in competition with, the homologous human transporter to modify the iron content of the phagosome environment[29, 42-44]. Iron is important for bacterial proliferation and competition for iron between host and pathogen has been well documented. Bacteria have previously been thought to evolve strategies for iron acquisition, further supporting the argument that *SLC11A1* could be a plausible candidate for coevolution between humans and MTB, although it has not been determined which MTB genes are involved[45-47]. *SLC11A1* polymorphisms have also been shown to associate with expression of IFN-γ, and the frequency of polymorphisms in *SLC11A1* are different between populations, including between Europeans and Africans[29, 42, 43]. Despite strong functional data indicating a biologically plausible role in the human response to TB, association studies of *SLC11A1* have shown inconsistent results with variants in this gene across global populations; analyses have shown that both the presence of statistical significance and the direction of effects vary between studies, both within and between populations[5, 48-60]. Specifically, an association between rs17235409, the specific SNP in the interaction term that we have replicated in both cohorts, and TB susceptibility has been previously published in five studies[50, 51, 56, 57, 59]. Of these five studies, two showed no association, one showed an association only among women below a certain age, and two showed a positive association with TB susceptibility. A recent study of susceptibility has also shown interaction between the Beijing strain of MTB and polymorphisms in *SLC11A1[29].* That *SLC11A1* has a strong biological justification but variable results in prior human genetic analyses make it a strong candidate for a gene susceptible to coevolution in which human association is affected by MTB lineage. Based on the current study, it is also possible that the inability to find a consistent association between *SLC11A1* genotype and TB disease could possibly be affected by the un-measured presence of different MTB lineages in prior studies that may have modified the association between *SLC11A1* and TB. While there may be other explanations for these findings, we think coevolution is the most likely reason for this interaction based on a well characterized function in the context of TB, the extent and variability of prior genetic epidemiological studies, and the fact that our statistical interaction results and predicted values are in line with previous literature studying coevolution.

## Materials and Methods

### Study participants

TB cases, extensive clinical data, and MTB isolates were collected as part of the Kawempe Community Health Study in Kampala, Uganda[61] (Supplemental methods). We examined two cohorts (N=113 and N=121, respectively), independently collected in Kampala. All TB cases were culture-confirmed; additional details about ascertainment and clinical characterization are provided elsewhere[61]. The two cohorts differed in percentage of HIV positive individuals (Table 1); therefore, HIV status was used as a covariate in all of our regression models. Previous analyses of microsatellite data from these cohorts indicated no substantial population substructure[62]. Symptoms utilized in the TBscore were evaluated upon diagnosis.

### Bandim TBscore

The TBscore is based on five self-reported symptoms: cough, hemoptysis, dyspnea, chest pain, and night sweats, as well as six signs identified at examination: anemia, pulse > 90 beats/min, positive findings at lung auscultation, temperature > 37 ° C, body mass index (BMI) < 18 kg/m^2^, and mid upper arm circumference (MUAC) < 220 mm. Each of the 11 clinical variables contributes 1 point, while BMI and MUAC contribute an extra point if <16 kg/m 2 and < 200 mm, respectively; thus, the maximum TBscore is 13.In our study, we did not have data on MUAC, so we instead used lean and fat mass body composition data obtained using bioelectrical impedance analysis (BIA), as described previously[63]. A TBscore of 8 or greater is associated with a 10% greater chance of death within 8 months than a TBscore under 8and a TBscore of 3 or higher is a significant predictor of mortality over 18 months of follow-up, even after adjusting for HIV status[32].

### Human genetic analysis

For our analysis of human host genetics, we examined 29 genes that affect innate and adaptive immune responses to TB chosen as part of a previous candidate gene study in Cohort 1. This candidate gene panel has been tested for association with TB but not with severity or TBscore[38]. Single nucleotide polymoprphisms (SNPs) in these genes were genotyped on a custom Illumina GoldenGate microarray. This analysis focused on genes in the Toll-like and Nod-like receptor families (*TLR1, TLR2, TLR4, TLR6, TLR9, TIRAP, TOLLIP, TICAM1/2, MyD88, NOD1, NOD2*), cytokines and their receptors expressed by macrophages *(TNF, TNFR1/2, IL1*α*/*β, *IL4, IL6, IL10, IL18, IL12A/B, IL12RB1/2, IFNG, IFNGR1/R2*), genes expressed by T-cells (*IFNG, IL4, IL12, STAT1, IL12RB1/2, IL10*) and key TB candidate genes (*SLC11A1, SLC6A3*). Haplotype tagging SNPs were selected to capture common genetic variation (minor allele frequency ≥ 5%) with strong coverage (linkage disequilibrium r^2^ ≥ 0.8) in any of the three African HapMap populations (YRI, LWK, MKK), based on previous analyses[64]. Many of these have previously been studied in animal, human, and macrophage models and are thought to be important in the human response to MTB infection[3-5, 65].

For Cohort 2, we used the Illumina HumanOmni5 microarray comprising 4,301,331 markers genome-wide, offering high genome wide coverage of common genetic variation even within African populations[66]. Genotype calling and quality control were performed as described in elsewhere[66]. Since the Cohort 2 data did not contain all the SNPs of interest from the Cohort 1 data, we used the Michigan Imputation server and protocols to impute SNPs[67, 68]. Low quality imputed SNPs (minimac r^2^ criterion < 0.5) were removed.

Only SNPs that had a call rate greater than 0.95 and MAF> 0.05 in both samples were used in the analysis. This resulted in a total of 403 eligible SNPs that were tested for association with TBscore and for interaction with MTB lineage (Supplemental Table 2) We examined all SNPs in Cohort 1, then attempted to replicate these in Cohort 2. As the two cohorts were similar to each other with respect to dependent and independent variables, we then combined the two cohorts into one and ran the analyses again on this combined cohort..

### MTB molecular analysis

MTB was isolated from sputum of each of these subjects and lineages were classified according to lineage-identifying SNPs using real-time PCR and validated with long sequence polymorphism PCR[18]. Lineage was determined from three SNPs that identify distinct lineages, as previously described by Gagneux[14, 16, 17]. Lineage 4 (L4) is the most geographically widespread and has historically been present across most of the world[14, 15]. L4 has many derived sub-lineages that each exist only within specific regions and to which people in these regions have been exposed for a shorter time[14, 15]. In the context of this study setting, the relevant MTB lineages were Lineage 4 (referred to in this paper as L4/Non-Ugandan), Lineage 3 (L3 also known as Central Asian), and Lineage 4.6/Ugandan, which is a specialist sub-lineage of L4 that is only found in Uganda and the countries immediately surrounding it (Supplemental Table 1)[14-18]. L4.6/Uganda is the most newly evolved of the three, a sub-lineage of the L4 generalist lineage, and is unique to this part of Africa[14, 15].

### Statistical analysis

The analysis of the marginal (main) effects analysis was done by analyzing SNPs as the independent variable and the TBscore as the dependent variable in a separate linear regression equation for each SNP, adjusting for HIV status as a binary covariate. SNPs were analyzed such that 1 was equal to the homozygous ancestral genotype and 0 was equal to the presence of the derived allele, as determined by the ENSEMBL and RefSeq reference genomes. We also analyzed these SNPs using ordinal regression and Poisson regression equations. Analysis was performed and figures generated using R version 3.5.2.

For our interaction term between lineage and human genotype, we chose to operationalize lineage as a binary variable. Each subject is coded as 1 for the L4.6/Ugandan lineage or as 0, which encompasses the L4/Non-Ugandan and L3/Central Asian lineages together. As both of these are older lineages and L4.6/Ugandan is a newer sub-lineage, this enables us to examine co-evolution as we are contrasting a lineage (L4.6) with which Ugandans have historically had shorter contact, relative to the two older lineages (L3 and L4) with which Ugandans have had longer contact. As we would expect a longer historical co-existence to associate with lesser severity and the introduction of a newer sub-lineage to associate with greater severity, we have grouped the two older lineages together. This also affords us greater power to detect a difference in our sample size than if we were to examine all 3 lineages independently.

We calculated a Bonferroni criterion for experiment-wide significance based on the number of SNPs analyzed that accounted for linkage disequilibrium (Supplemental methods).

## Acknowledgements

Funding for this work was provided by the Tuberculosis Research Unit (grant N01-AI95383 and HHSN266200700022C/ N01-AI70022 from the NIAID) and R56 AI130947.We would like to acknowledge the invaluable contribution made by the study medical officers, health visitors, laboratory and data personnel: Dr. Lorna Nshuti, Dr. Roy Mugerwa, Dr. Sarah Zalwango, Dr. Chriospher Whalen, LaShaunda Malone, Dr. DeoMulindwa, Dr. Christina Lancioni, Allan Chiunda, Denise Johnson, Bonnie Thiel, Mark Breda, Dennis Dobbs, Hussein Kisingo, Mary Rutaro, Albert Muganda, Richard Bamuhimbisa, Yusuf Mulumba, Deborah Nsamba, Barbara Kyeyune, Faith Kintu, Mary Nsereko, Gladys Mpalanyi, Janet Mukose, Grace Tumusiime, Pierre Peters, AnnetKawuma, SaidahMenya, Joan Nassuna, Alphonse Okwera, Keith Chervenak, Karen Morgan, Alfred Etwom, Micheal Angel Mugerwa, and Lisa Kucharski. We would like to acknowledge and thank Dr. Francis AdatuEngwau, Head of the Uganda National Tuberculosis and Leprosy Program, for his support of this project. We would like to acknowledge the medical officers, nurses and counselors at the National Tuberculosis Treatment Centre, Mulago Hospital, the Ugandan National Tuberculosis and Leprosy Program and the Uganda Tuberculosis Investigation Bacteriological Unit, Wandegeya, for their contributions to this study. This study would not be possible without the generous participation of the Ugandan patients and families.

## Description of Supplemental Materials

We have provided: a pdf document with supplemental text on methods including 1 supplemental table and 3 supplemental figures, and 2 supplemental tables in the form of Microsoft Excel spreadsheets.

