## Supplemental Text for "Interaction between host genes and M. tuberculosis lineage can affect tuberculosis severity: evidence for coevolution"

**Study Population**

The study population includes participants from two independently recruited cohorts in Ugandan households. The first cohort of subjects, which will be referred to as Cohort 1, were enrolled in a household contact study that enrolled patients from 1995 to 1999 and from 2002 to 2012[1, 2]. Cohort 2, which is an independent cohort of subjects, were enrolled between 2002-2012. The original study protocol was reviewed and approved by the National HIV/AIDS Research Committee, The Uganda National Council of Science and Technology and the institutional review board at the University Hospitals Case Medical Center, Cleveland, OH, USA. Patients who presented to the study clinic with active pulmonary TB were enrolled as index cases. Diagnosis of pulmonary TB for the present study was confirmed based on isolation of MTB from clinical gastric or sputum samples. A total of 141 index cases from Cohort 1 and 135 from Cohort 2 met these criteria. We then excluded 19 cases from Cohort 1 and 9 from Cohort 2 because these participants were under 15 years old and our outcome measure was not valid to assess TB severity in patients this young. We further excluded 7 cases from the Cohort 1 and 3 from Cohort 2 who had missing information for the MTBlineage. After these exclusions there were 115 cases fromCohort 1 and 123 cases from Cohort 2 but two people were in both cohorts and were removed from both leaving 113 and 121 for analyses.

**Clinical Phenotyping**

The BandimTBscore is based on five self-reported symptoms: cough, hemoptysis, dyspnea, chest pain, and night sweats, as well as six signs identified at examination: anemia, pulse > 90 beats/min, positive findings at lung auscultation, temperature > 37 ° C, body mass index (BMI) < 18 kg/m^2^ , and mid upper arm circumference (MUAC) < 220 mm. Each of the 11 clinical variables contributes 1 point, while BMI and MUAC contribute an extra point if <16 kg/m 2 and < 200 mm, respectively; thus, the maximum score is 13[3-5]. In our study, we did not have data on MUAC, so we instead used lean and fat mass body composition data obtained using bioelectrical impedance analysis (BIA), as described elsewhere[6-8].

**Genotyping**

For Cohort 1, a custom Illumina GoldenGate 10k microarray was designed for a previous analysis of TB candidate genes[9]. This analysis focused on genes in the Toll-like and Nod-like receptor families (*TLR1, TLR2, TLR4, TLR6, TLR9, TIRAP, TOLLIP, TICAM1/2, MyD88, NOD1, NOD2*), cytokines and their receptors expressed by macrophages *(TNF, TNFR1/2, IL1α/β, IL4, IL6, IL10, IL18, IL12A/B, IL12RB1/2, IFNG, IFNGR1/R2*), genes expressed by T-cells (*IFNG, IL4, IL12, STAT1, IL12RB1/2, IL10*) and key TB candidate genes (*SLC11A1, SLC6A3*). Many of these have previously been studied in animal, human, and macrophage models and are thought to be important in the human response to MTB infection[10-13].

Haplotype tagging SNPs were selected to capture common genetic variation (minor allele frequency ≥ 5%) with strong coverage (linkage disequilibrium r^2^ ≥ 0.8) in any of the three African HapMap populations (YRI, LWK, MKK), based on previous analyses[14]. Tag SNPs were identified using Genome Variation Server (GVS) (http://gvs.gs.washington.edu/GVS137/index.jsp). Genotyping was conducted using the Illumina iSelect platform. Once SNPs were selected using GVS, their availability on the iSelect platform was verified; if a specific SNP was not available on iSelect, a nearby SNP was selected to replace it. Genotype calling and quality control was performed using Genome Studio, filtering the SNPs by call frequency, replicate errors, and clustering quality. Family relationships were corrected and resolved where needed, including defining subfamilies of first-degree relatives within households.

For Cohort 2, we used the Illumina HumanOmni5 microarray comprising 4,301,331 markers genome wide,offering high genome wide coverage of common genetic variation even within African populations[15]. Genotype calling and quality control were performed as described in a previous publication[15].

**Imputation**

Since theCohort 2 data cohort did not contain all the SNPs of interest from the Cohort 1 data, we used the Michigan Imputation server and protocols to impute SNPs[16, 17]. Low quality imputed SNPs (minimac r^2^ criterion < 0.5) were removed. Only SNPs that had a call rate greater than 0.95 in both samples were used in the analysis. This resulted in a total of 403 eligible SNPs. For the marginal analyses, 3 of the 403 SNPs returned an error message. For the interaction analyses in Cohort 1, 3 of 403 SNPs returned an error. In Cohort 2,8 SNPs returned an error. 393 SNPs have results for the interaction analyses in both data cohorts (did not return any error messages). These errors results from a MAF<0.05.

**Lineages**

Mycobacterium tuberculosis (MTB) has distinct genetic lineages historically associated with different regions of the world. MTB isolates were obtained from study participants and real time PCR was performed to genotype three SNPs used to classify MTB into lineages, as previously described by Gagneux et al. [18, 19]. This sample of Ugandan subjects comprised three lineages: L4.6/Uganda, L4/Non-Ugandan, and L3-Central Asian, as described in the main text.

**Statistical Analysis**

All analysis was performed using R version 3.5.2. The association between SNPs and the Bandim TBscore and the interaction between SNPs and MTB lineage was assessed using linear regression models. We assessed the relationship between SNPs and TB severity on the Bandim TB scale. Each SNP was the independent variable in a separate regression equation and were coded using a dominant model. The analysis included HIV status as a covariate. Nominal significance was defined as p<0.05, and replication was based on the same SNP having p<0.05 with effect in the same direction in both cohorts.

We analyzed the interaction between SNP and Ugandan lineage by creating multiplicative interaction terms for SNP and Ugandan lineage in a linear regression model. All analysis was done in R. This analysis also included HIV status as a covariate. Because of small sample sizes we did not perform similar analyses for the L4/Non-Ugandan or L3:Central Asian lineages. All SNPs from Cohort 1 were analyzed in Cohort 2. P-values below 0.05 for the interaction term in both datasets with coefficients in the same direction in both cohorts (both positive or both negative) were considered to be significant results. We then ran a combined analysis to generate meta P-values and the threshold for significance was corrected using a Bonferroni correction that is adjusted for linkage disequilibrium (LD) within the dataset by finding the equivalent number of independent tests of association. This adjustment is necessary for SNPs that are imputed using LD structure as many of the tests being run are correlated[20]. The Bonferroni correction yielded a threshold of p=0.00023 for our 403 SNPs, equivalent to 371 independent tests. For our predictions of the count of symptoms, we created a Poisson model of the interaction between rs17235409 and lineage that adjusted for HIV status as a covariate. We then used the predict function in R and rounded the predictions to generate counts of symptoms for each of the four combinations of lineage and rs17235409 genotype (Supplemental Table 4).

**Supplemental Table 1. Distribution of Lineages in each cohort**

|  | Cohort 1 | Cohort 2 | Total | p |
| --- | --- | --- | --- | --- |
| L4-Ugandan | 73 (64.6) | 73 (60.3) | 146 (62.4) |  |
| L4-Non-Ugandan | 29 (25.7) | 31 (25.6) | 60 (25.6) |  |
| L3-Central Asian | 11 (9.7) | 17 (14.0) | 28 (12.0) |  |
| Total | 113 | 121 | 234 | 0.58 |

Figures are shown as N (%). P-value is from a χ^2^ test of independence

**Supplemental Table 4. Rounded Count of Symptoms from Poisson Model Predictions**

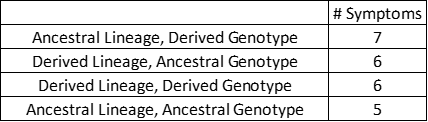

**Supplemental Figure 1. Haploview LD Plot for IL12B in Cohort 1**

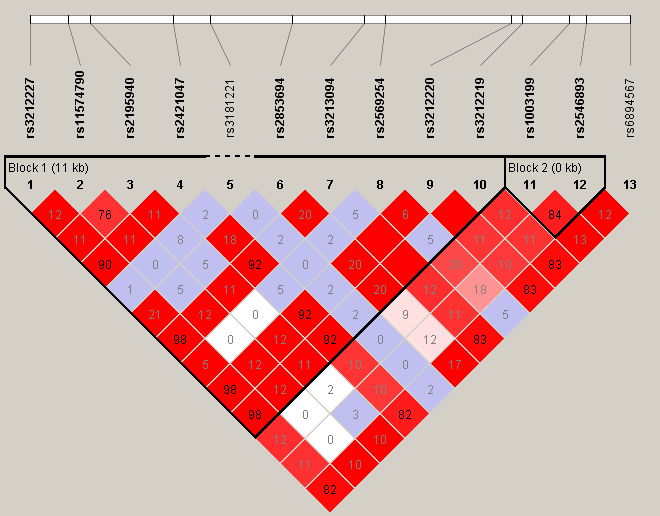

**Supplemental Figure 2. Haploview LD Plot for IL12B in Cohort 2**

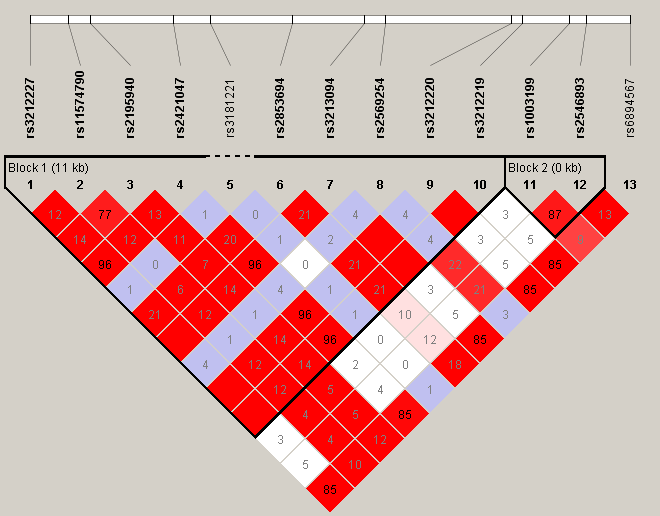

**Supplemental Figure 3. Haploview LD plot for IL12B in Combined Dataset**

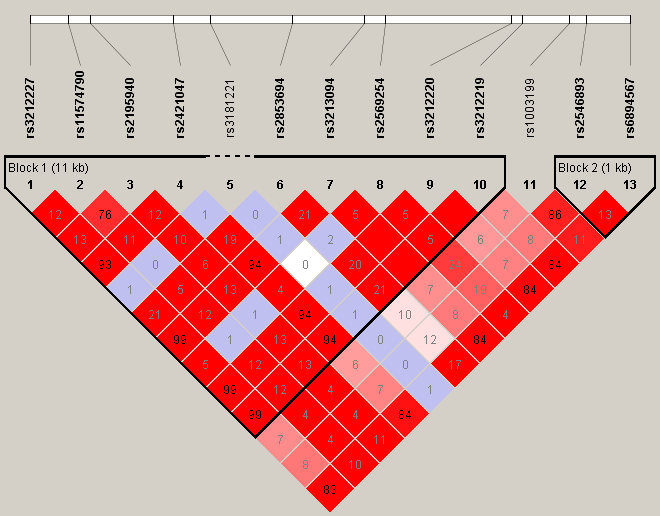
